# ConTreeDP: A consensus method of tumor trees based on maximum directed partition support problem

**DOI:** 10.1101/2021.10.13.463978

**Authors:** Xuecong Fu, Russell Schwartz

## Abstract

Phylogenetic inference has become a crucial tool for interpreting cancer genomic data, but continuing advances in our understanding of somatic mutability in cancer, genomic technologies for profiling it, and the scale of data available have created a persistent need for new algorithms able to deal with these challenges. One particular need has been for new forms of consensus tree algorithms, which present special challenges in the cancer space for dealing with heterogeneous data, short evolutionary time scales, and rapid mutation by a wide variety of somatic mutability mechanisms. We develop a new consensus tree method for clonal phylogenetics, ConTreeDP, based on a formulation of the Maximum Directed Partition Support Consensus Tree (MDPSCT) problem. We demonstrate theoretically and empirically that our approach can efficiently and accurately compute clonal consensus trees from cancer genomic data.

Availability: https://github.com/CMUSchwartzLab/ConTreeDP

## 1 Introduction

Tumor progression has long been recognized as a process of clonal evolution [1], in which tumors develop through the accumulation of genetic diversity in clonal cell lineages and selection for variants that tend to promote tumor growth, migration, escape from treatment, or other phenotypes of cancer progression. This recognition led to the development of cancer phylogenetics [2], the use of phylogenetic algorithms to reconstruct the mutational history of cell lineages in a cancer, typically depicted as rooted trees with nodes labelled via unique mutations during tumor progression. These “tumor phylogenies” provide a way to characterize tumor heterogeneity, compare tumor genesis and progression among different patients with the same cancer type, or collectively between different cancer types, and provide insights into the forces of evolutionary drift and selection that drive tumor growth. With the explosive growth in the size and complexity of cancer genomic data sets, especially as single-cell genomics has become prominent in cancer studies [3, 4], tumor phylogenetics has become an indispensible tool for interpreting tumor genomic data. For reviews of the field, see [5].

Many computational methods have now been developed for inferring clonal evolutionary trees for various data types (e.g., single-cell, bulk whole genome, or targeted deep sequencing) and measures of genetic or epigenetic diversity (e.g., copy number alterations (CNAs), single nucleotide variants (SNVs), or methylation profiling). While different data types and algorithms can yield complementary insights into tumor evolution, interpretation can be challenging when different techniques yield distinct trees on a common set of samples. Similarly, distinct algorithms or chance sequencing errors may yield different results on a single data set. Replicate solutions from stochastic approaches like the common Markov chain Monte Carlo (MCMC) sampling [6, 7, 8] or combinatorial optimization methods [9, 10] or combinatorial enumeration methods like [11, 12] can also lead to multiple trees for a single data set [13, 14, 5]. Each of these scenarios leads to the challenge of how best to integrate the inferred evolutionary information from different trees to yield more detailed, accurate, and comprehensive mutation histories.

The general problem of reconciling a collection of distinct trees describing a common set of taxa is known as a consensus tree problem, and development of methods for this problem has long been a concern of the phylogenetics community. Numerous consensus tree methods have been developed for species phylogenetic inference based on different criteria (see [15] for a review). One of the major classes is quartet-based phylogenetic methods that typically solve a version of the Maximum Quartet Consistency Problem (MQC) to identify a phylogeny that maximizes number of quartets in a global quartet set usually derived from a set of trees. One particularly influential recent method has been ASTRAL [16], a dynamic programming method for efficient consensus tree inference scalable to whole genome datasets from large numbers of species.

Tumor phylogenetics shares similar challenges to species phylogenetics but also some important differences, leading more recently to a literature of specialized consensus methods specifically for tumor phylogenies. For example, in species phylogenetics it is typically assumed that all leaves of a tree are labeled with the species and internal nodes represent unobserved ancestral states, while tumor evolutionary histories often involve mixed populations of both ancestral and leaf nodes [13]. The chronological history of mutations is also important as it is more comparable among different patients or types of cancer than the clonal evolution history. Such challenges led [13] to develop four simple tree distance metrics and an associated consensus tree method, GraPhyC, specifically for tumor evolution. Based on that work, [17] then developed a multiple consensus tree problem to solve for tree clustering. While these methods represent an important advance for tumor phylogenetics, there is no current method that can obtain reliable consensus tree particular for complicated combinations of mutation types. For example, GraPhyC requires a preclustering step to identify node sets that can obscure some true variation and can introduce error when dealing with important genomic variant types such as CNAs and SVs that require specialized phylogeny methods [18].

Here, we pose the consensus problem as a maximum directed partition problem, where we maximize the directed partitions in rooted directed mutation tree, similar to the Maximum Quartet Consistency Problem (MQC) from species phylogenetics. We use this model to formulate a new consensus tree method for tumor phylogenetics, using techniques similar to those used to solve MQC by the widely used ASTRAL method [16], to allow for improved generality with respect to the sources of the input trees. Specifically, we pose a maximum directed partition support consensus tree (MDPSCT) problem to directed labeled tree with the nodes single-labeled or multilabeled with mutations from a total set of mutations. This model leads to our method ConTreeDP, a dynamic programming method to achieve a polynomial algorithm given a maximum allowed node degree for the consensus tree. ConTreeDP can identify an optimal tree without requiring that it appear in the input trees or the optimal clustered mutation labels, allowing more fine-scaled resolution of mutation histories than prior work. The method also allows for an adjustable granularity for resolving or preserving uncertainty due to low-support branches. We validate the method on simulation data, showing improved performance in reconstructing the true history compared to GraPhyC. We further apply ConTreeDP to the same real data as examined in [14] yielding corrections to inferred mutation histories that we argue are more likely to reflect the true biology.

## 2 Method

### 2.1 Problem formulation

In species phylogeny, quartet based methods are among the category of consensus methods that infer the evolutionary tree based on substructures derived from a given set of trees. The Maximum Quartet Consistency (MQC) problem is an intuitive problem to maximize the number of observed quartets, based on which many efficient algorithms have been developed. Due to the distinct properties of tumor trees from species trees, including multi-labeled nodes and labeled internal nodes, we propose a modification of that idea that we call the Maximum Direct Partition Consistency (MDPC) problem, which maximizes the number of directed partition topologies derived from a given set of tumor trees. We define the directed partition as follows:

#### Definition

For a directed tumor tree *T* with labels set *S*, we define a **directed partition** 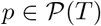 corresponding to a directed branch *b* = (*u*, *v*) ϵ *T* as (*S*_1_, *S*_2_) where *u*, *v* are parent and child pair of nodes with multiple labels, *S*_2_ = {*x*|*x* ϵ *L*(*w*), *w* ϵ *D*(*v*)} + {*L*(*v*)} and *S*_1_ = *S* \ *S*_2_, where *D*(*v*) is the set of nodes descended from *v* in *T*, *L*(*w*) is the set of labels labeled at node *w*.

Note that because *T* is a tree, *S*_1_ and *S*_2_ become disconnected in *T* if *b* is removed.

The goal of ConTreeDP is to derive a consensus tumor tree from a set of tumor evolutionary trees generated from different methods, different replicates of a single method, or even different data sets. As with ASTRAL, we formulate this informally as an optimization problem of seeking a tree structure whose edge set (defined as a set of directed partitions) has the highest total support in a set of rooted input trees given a set of allowed directed partitions. Intuitively, mutations that occur nearby on most input trees should preferentially be grouped or close in the consensus tree, based on the assumption that there will be some but not many discrepancies between the tree structures of the input trees.

We formulate the goal of our method as solving the Maximum Directed Partition Support Consensus Tree (MDPSCT) problem (Fig.1):

**Input**: A set of mutation labels *C*, a set of directed rooted tumor trees 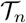 where each node *v_n_* in each *T_n_* is labeled with one or more mutation labels set *c* ⊆ *C*, and a directed partition set Δ where each *δ* ϵ Δ can be characterized as a directed partition of *C*.

**Output**: A directed tumor consensus tree *T* such that 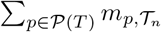 is maximized and 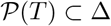, where 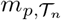 is the number of occurrences of directed partition *p* in tree set 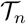

**Figure 1:**
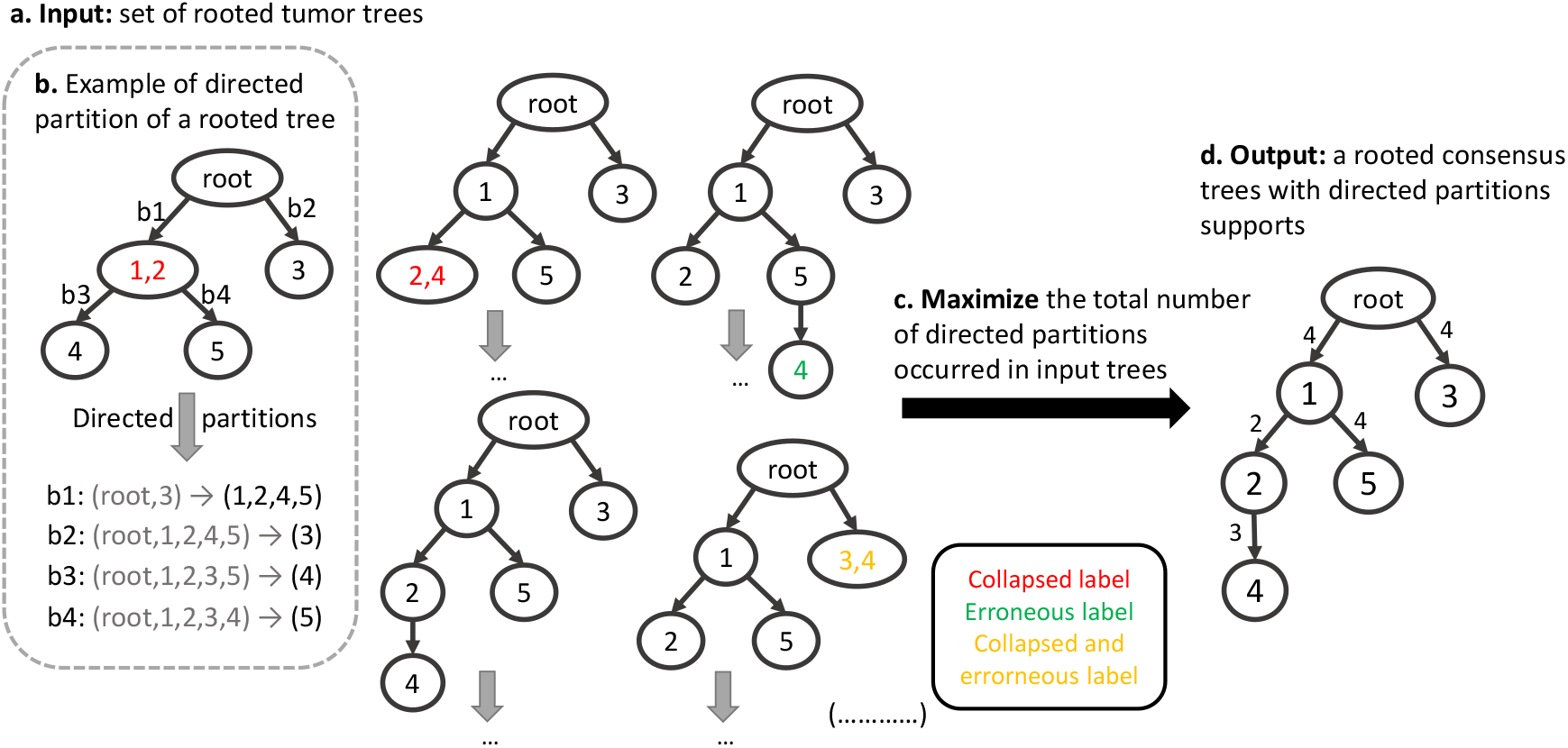
Illustration of the MDPSCT problem. a. Example input data set consisting of a collection of rooted tumor trees. b. Illustration of directed partitions of a rooted tree, each of which can be characterized by the set of labels labeling the nodes in the sink set of the partition. c. Statement of the objective function. d. An optimal output tree for the sample input with each edge labeled by the supported of its corresponding directed partition.

If Δ is the set of all possible directed partitions of the labels set *C*, then the solution to the problem will be the true maximum support consensus tree. As with the ASTRAL method [16], runtime is dependent on |Δ|, however, so to derive an efficient algorithm we restrict the solution to a smaller set observed in the input data. In practice, we define 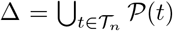 to be the set of all directed partitions occurring in our input tree set 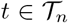, which leads to a polynomial time algorithm for the problem given a fixed maximum node degree.

### 2.2 Approach

The core of our approach for solving the MDPSCT problem with a fixed mutations set is a dynamic programming (DP) method, which is a similar algorithmic strategy to ASTRAL despite the differences in the models. Fig. 2 shows a visualization of the method with a toy example. We assume that in preprocessing, the total number of occurrences of each possible directed partition in the input tree set has been counted. The dynamic programming algorithm then recursively finds the best possible directed partition support and corresponding subtree structure for a given set of labels *s* which belongs to the sink set partitions of the given directed partition set. The structure of the full tree can then be obtained by backtracking on the optimal subset combinations built through the dynamic programming. Optionally, the detailed tree can be collapsed on the branches with low support based on a pre-defined threshold. We specify a maximum degree *d*, which is a maximum number of children of any parent node allowed in the output tree. The result is an algorithm whose runtime is provably polynomial for fixed *d*.

**Figure 2:**
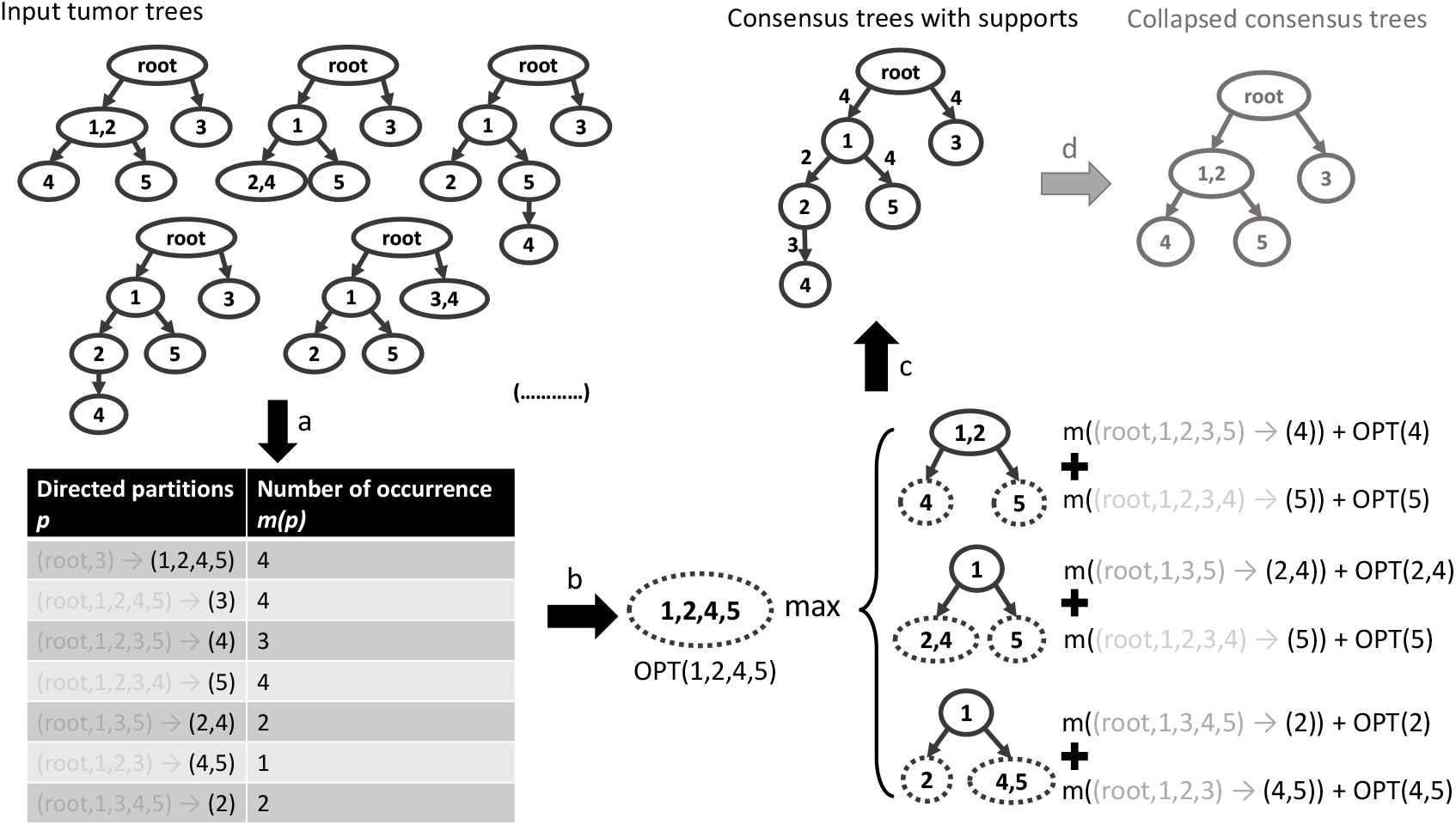
Visualization of the ConTreeDP method with a toy example. a. Summarizing the total count of all directed partitions (characterized as sink set partitions) in the input trees 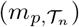. b. Identifying the optimal support for each possible subset of mutations. We demonstrate the process for recursively calculating the optimal support for the subset of (1,2,4,5). c. Back tracking the optimal tree structure to identify the optimal directed partition supports. d. (Optional) collapsing branches with low support based on a pre-defined threshold.

We define the optimal support of a subset of mutation labels *s* ⊂ *C* as opt(*s*). Therefore, any possible subsets 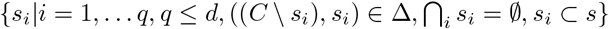 can consist of s with the element 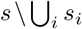 being the relative ancestor node of the subset *s*. The optimal support of *s* consists of the total optimal support of *s_i_* for all *i* and the total counts for directed partitions (*C* \ *s_i_*, *s_i_*) for all *s_i_*. Therefore the recursion formula can be expressed as follows:

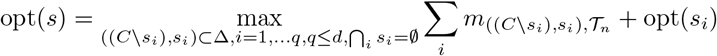

At each step of the recursion, then, we need to partition the nodes of a rooted subtree in all possible ways consistent with the allowed set of directed partitions Δ and the maximum node degree *d* before selecting the partition that yields maximum support for the given subtree.

### 2.3 Properties

#### 2.3.1 Runtime analysis

The runtime for the algorithm is *O*((*nk*)^*d*^). For the preprocessing step, the runtime for calculating the total count for each directed partition observed in all input trees is *O*(*nk*). In the dynamic programming step, we traverse all possible combinations of mutations with numbers of set from 1 to *d* for each observed partition, resulting in a runtime of *O*((*nk*)^*d*^). The final backward traversal, to obtain the optimal tree structure, takes *O*(*k*) time. Therefore the total runtime is *O*((*nk*)^*d*^). In real practice, *d* might be set to the largest degree found in any input tree, or set heuristically based on resource limits or prior expectations.

#### 2.3.2 Consistent estimator

The solution to our problem yields a consistent estimator of the directed tree.

Next we want to prove why it is reasonable to use this objective.

##### Proof Sketch

According to parsimony principle, it is intuitive that the probability of identifying a certain partition defined by a branch *b* = (*u*, *v*) in a mutation history tree *t* is

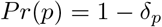

where *δ_p_* is the error rate for mistakenly identifying another incompatible partition and where we assume *Pr*(*p*) > *δ_p_*.

We define a set of random variables *X*_*pt*1_, *X*_*pt*2_… *X_ptn_* ~ Bernoulli(*Pr*(*p*)) as 0-1 variables, indicating whether tumor tree *t*_1_, *t*_2_…*t_n_* identify a particular partition *p*. We assume these variables are i.i.d. Therefore, we define 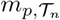 as the number of trees in 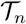 that identify the partition *p*.

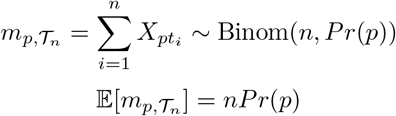

Since we know that 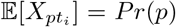, Var(*X_pti_*) = *Pr*(*p*)(1 – *Pr*(*p*)) < ∞, according to the Weak Law of Large Numbers:

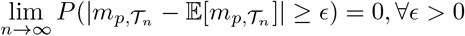

Then we want to show that the true tree 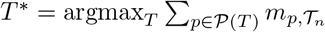 when *n* is large for the exact case. Assume that *T′* is a rooted tree other than *T*\*. Therefore, there exists at least one partition 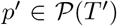 but 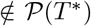. Symmetrically, there at least exists one partition 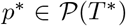 but 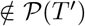.

Since we generate the tree set 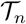 from data derived from an underlying true tree, the probability of identifying partition *p′* is some *δ* < *δ_p_*.

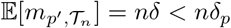

Similarly,

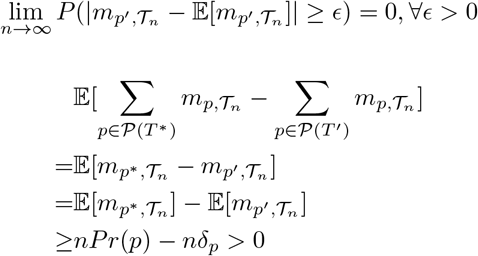

According to Slutsky’s theorum:

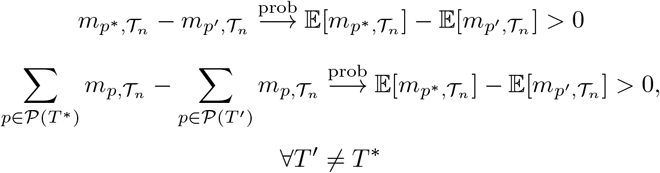

Therefore the method is consistent in estimating the optimal solution. The method also solves the problem when it restricts the tree search to an induced directed partition set 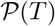 that is a subset of the Δ, the set of all directed partitions occurring in the tumor trees 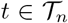. When *n* is large, from WLLN, 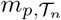 will be close to *nPr*(*p*) for 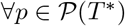, therefore 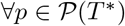 will occur in 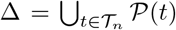 with high probability, which means that subject to the restriction to Δ, the method will also find the optimal solution.

## 3 Results

### 3.1 Simulation data

For each trial, we randomly simulate a tumor tree and generate a series of perturbed trees by applying a process of randomly collapsing edges and a fixed rate of randomly moving subtrees, as was done by [14]. To be more specific, we first randomly generate a mutation tree topology with a maximum degree *d* =3 with a given number of nodes *k* as the ground truth, where each node represents a mutation. We also assign a frequency to each node so that it obeys the sum rule, i.e., that the frequency of any ancestor mutation should be higher or equal to the sum of the frequencies of its children[8]. We then randomly collapse the parent-child pairs with probability of 0.9 if the difference between the frequency of a parent mutation and the sum of frequency of the child mutations is smaller than 0.01, with probability of 0.5 if the difference if smaller than 0.05, and with probability of 0.01 otherwise. Then we randomly move subtrees to other parent nodes consistent with the sum rule with probability of 0.1 for each branch.

We evaluate ConTreeDP based on its ability to accurately reconstruct the original trees. We compare ConTreeDP’s performance with that of GraPhyC [13]. We use the CASet and DISC distance measures introduced in [19] for evaluating the distance between the inferred tumor trees and the ground truth tree. CASet emphasizes the common ancestors of mutations so as to penalize more those changes close to the root of the tree, while DISC relies more on distinctly inherited ancestors, resulting in more emphasis on switches among more recent mutations. These distance measures were specifically developed for tumor mutation trees and account for some of their unique inheritance properties, most importantly by taking internal nodes into comparison, unlike traditional phylogenetic distance metrics like the Roubinson-Foulds (RF) distance.

Overall, our method yields improved accuracy in inferring consensus trees compared with GraPhyC in various settings (Fig.3). There are few more outliers with higher mutations number when using the CASet measure compared with GraPhyC, especially when number of trees is small. This results from the random movement of subtrees with a relatively high fixed rate when the mutation number is large, causing drastic changes in tree structures in ways that might not accurately reflect real data.

**Figure 3:**
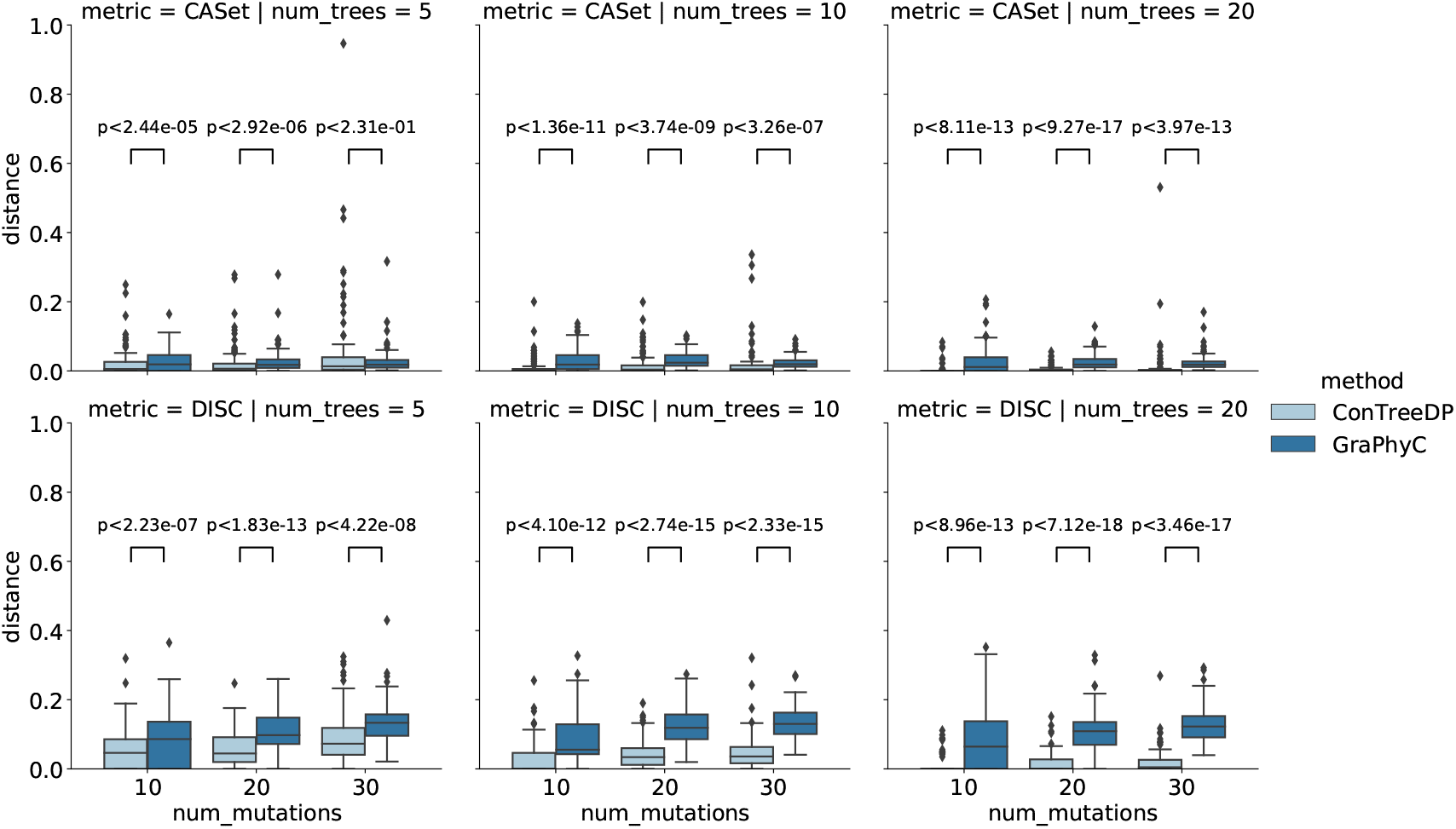
The simulation result with different parameter settings. We scanned input tree number 5, 10 and 20, number of mutation 10, 20 and 30. Distance measures CASet and DISC were used for evaluation. Wilcoxon rank sum test is used for testing. Each parameter configurations has 100 repeats.

### 3.2 Real data

We applied our method to two real cases used in GraPhyC validation [13, 14]: a chronic lympocytic leukemia (CLL) and a triple negative breast cancer (TNBC) case.

#### 3.2.1 CLL

Chronic lymphocytic leukemia sample CLL077 from [20] has been widely used as a benchmark dataset for prior deconvolutional methods. Among these, PhyloWGS [6] was used with multiple random starts by [13, 14] to validate GraPhyC on real data. We used the same CLL input trees as [13, 14] and applied ConTreeDP to these eight tumor trees (Fig.4(a)(b)). The weight label on each branch shows the support of the directed partition corresponding to that edge. GraPhyC preclusters mutation labels while ConTreeDP does not, yielding potentially different granularities of solutions. In order to allow a fair comparison between GraPhyC and ConTreeDP, then, we added to ConTreeDP a post-processing step to collapse those branches with length support smaller than half of the total number of input trees. In this case, branches with support lower than 4 are collapsed. We can see that the collapsed consensus tree from ConTreeDP is identical to the consensus tree from GraPhyC with all four distance measures, showing the ability of ConTreeDP to produce a reliable consensus tree while retaining fine granularity and producing a more detailed mutation history (Fig.4(c)(d)).

**Figure 4:**
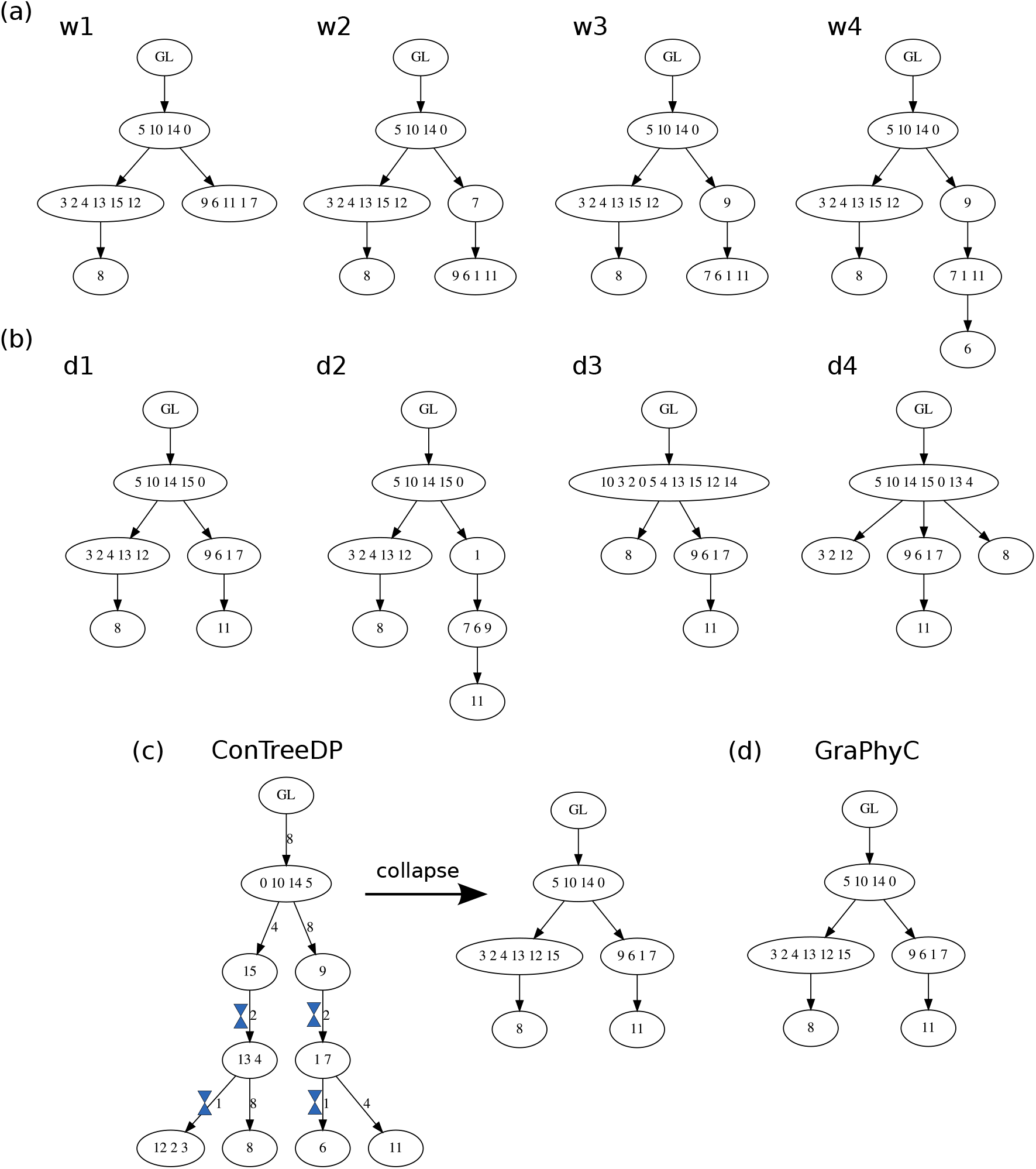
The input trees and output results of ConTreeDP and GraPhyC on CLL077. (a) Four input trees obtained from PhyloWGS using whole-genome sequencing of CLL077. (b) Four input trees obtained from PhyloWGS using ultra deep sequencing. (c) The consensus tree obtained by ConTreeDP and corresponding tree after collapsing the branch with support smaller than half of the total tree. (d) The consensus tree obtained by GraPhyC

#### 3.2.2 TNBC

[21] exome-sequenced 16 single tumor nuclei and 16 normal tumor nuclei of triple-negative breast cancer. [22] used the dataset to generate three tumor evolutionary trees by distinct inference methods: PhISCS[23], SiFIT[24] and SCITE[25]. [14] used the trees as input to GraPhyC to obtain a consensus tumor tree. We applied ConTreeDP to the same input trees and compared with the consensus tree inferred by GraPhyC. We also collapsed the tree from ConTreeDP with the same criterion as with the CLL dataset for fairer comparison.

The trees are very similar but with one difference, corresponding to mutation MAP3K4. All consensus trees by GraPhyC across different measures inferred MAP3K4 mutation (marked by blue rectangular frames) to occur relatively late, although with different positions or trajectories across trees (Fig.5(e)). By contrast, the ConTreeDP consensus tree inferred MAP3K4 to be a relatively early mutation in the tumor progression (Fig.5(d)). Two of the three input trees, those using PhISCS (Fig.5(a)) and SCITE (Fig.5(c), inferred MAP3K4 to be an early mutation, whereas the tree from SiFIT inferred the opposite (Fig.5(b)). We thus believe the ConTreeDP result to be more plausible than those derived from GraPhyC on this case.

**Figure 5:**
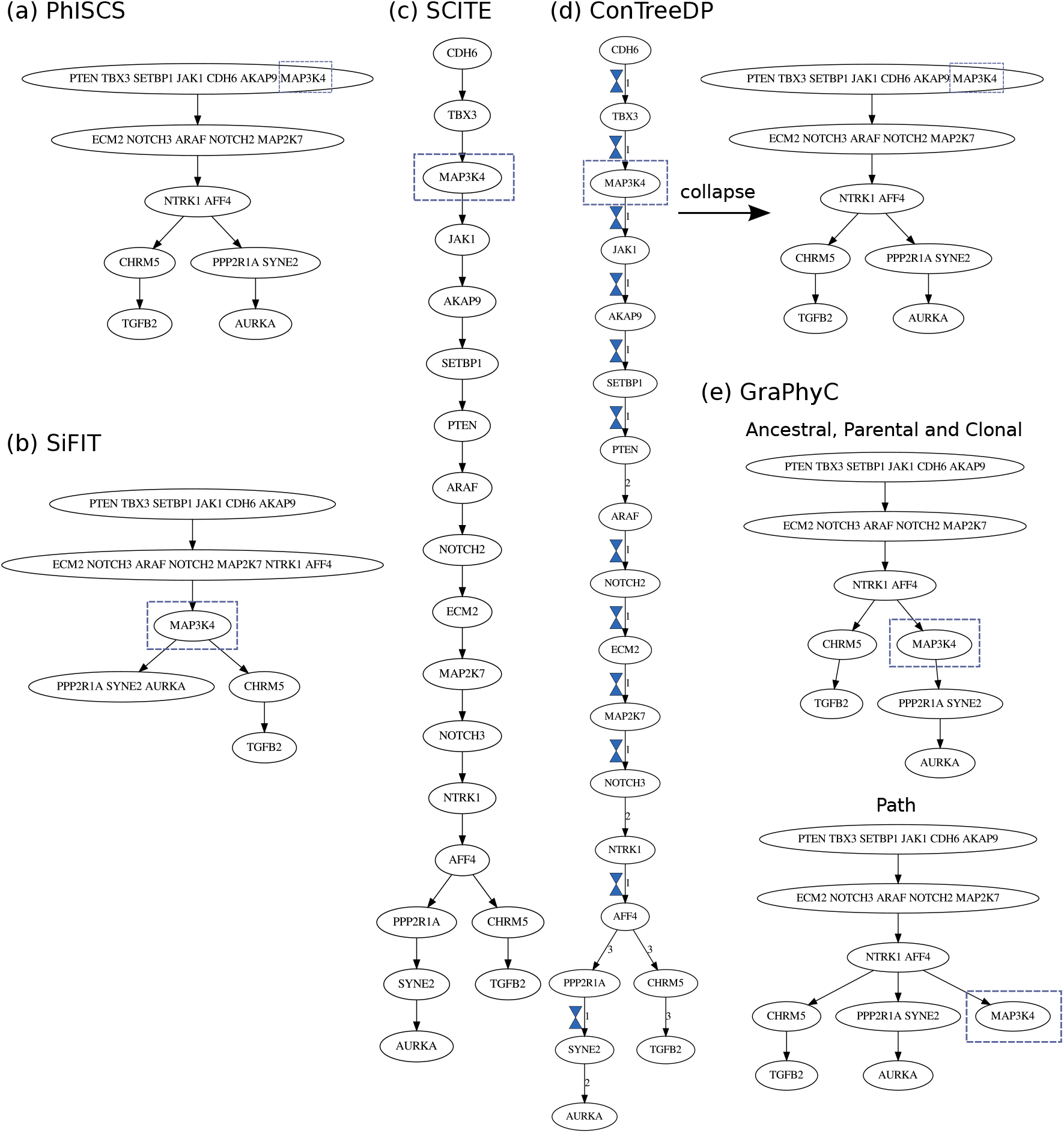
Input tree obtained from (a) PhISCS, (b) SiFIT, (c) SCITE, (d) The consensus tree obtained from ConTreeDP and corresponding tree after collapsing the branch with support smaller than half of the total tree. (d) The consensus tree obtained by GraPhyC

## 4 Discussion

We introduced a new MDPSCT problem formulation for deriving a consensus tree from multiple tumor history trees and developed an algorithm, ConTreeDP for efficiently identifying optimal trees by this model. We proved the method yields a consistent estimator and a polynomial time algorithm when solutions are restricted by a maximum node degree and to using observed directed partitions. We evaluated ConTreeDP on simulated data and showed better performance compared with a current state-of-the-art alternative. We also applied it to two real data sets and showed reliable consensus tree results that are consistent with prior work and show at least some likely improvement. The results suggest ConTreeDP can be a valuable addition to tools for building tumor evolutionary histories and other forms of cell lineage trees.

The method in its current form does have limitations, however. Most notably, it requires the input mutation labels to be the complete set of mutations for all trees. We work around this limitation currently by removing those mutations do not exist in all trees, a solution that might not generalize well to larger or noisier datasets. Finding better solutions could help the method generalize to more large datasets and help bring it to other applications, such as integrating trees from heterogeneous data sources or acting as a summary method for subsampled or other incomplete genomic data sets.

## Acknowledgment

We would like to thank Yifeng Tao for giving feedback, and also thank Haoyun Lei and Arjun Srivatsa for their assistance. Portions of this work have been funded by US NIH award R21CA216452 and Pennsylvania Dept. of Health awards 4100070287 and FP00003273. Research reported in this publication was supported by the National Human Genome Research Institute of the National Institutes of Health under award number R01HG010589. This work was also partially supported by an AWS Machine Learning Research Award. The content is solely the responsibility of the authors and does not necessarily represent the official views of the National Institutes of Health. The Pennsylvania Department of Health specifically disclaims responsibility for any analyses, interpretations or conclusions.

## Notes

### Competing Interest Statement

The authors have declared no competing interest.

